# Detection of YAP1 and AR-V7 mRNA for Prostate Cancer prognosis using an ISFET Lab-On-Chip platform

**DOI:** 10.1101/2022.08.04.502773

**Authors:** Joseph Broomfield, Melpomeni Kalofonou, Thomas Pataillot-Meakin, Sue M. Powell, Nicolas Moser, Charlotte L. Bevan, Pantelis Georgiou

**Affiliations:** Centre for Bio-Inspired Technology, Department of Electrical and Electronic Engineering, Imperial College London, London, SW7 2AZ, United Kingdom; Imperial Centre for Translational and Experimental Medicine, Department of Surgery and Cancer, Imperial College London, London, W12 0NN, United Kingdom; Sir Michael Uren Hub, Department of Bioengineering, Imperial College London, London, W12 0BZ, United Kingdom; Molecular Science Research Hub, Department of Chemistry, Imperial College London, London, W12 0BZ, United Kingdom

## Abstract

Prostate cancer (PCa) is the second most common cause of male cancer-related death worldwide. The gold standard of treatment for advanced PCa is androgen deprivation therapy (ADT). However, eventual failure of ADT is common and leads to lethal metastatic castration resistant PCa (mCRPC). As such, the detection of relevant biomarkers in the blood for drug resistance in mCRPC patients could lead to personalized treatment options. mRNA detection is often limited by the low specificity of qPCR assays which are restricted to specialised laboratories. Here, we present a novel reversetranscription loop-mediated isothermal amplification (RT-LAMP) assay and have demonstrated its capability for sensitive detection of AR-V7 and YAP1 RNA (3×10^1^ RNA copies per reaction). This work presents a foundation for the detection of circulating mRNA in PCa on a non-invasive Lab-on-chip (LoC) device for use at point-of-care. This technique was implemented onto a Lab-on-Chip platform integrating an array of chemical sensors (ion-sensitive field-effect transistors - ISFETs) for real-time detection of RNA. Detection of RNA presence was achieved through the translation of chemical signals into electrical readouts. Validation of this technique was conducted with rapid detection (*<*15 min) of extracted RNA from prostate cancer cell lines 22Rv1s and DU145s.

## Introduction

One in eight men are expected to be diagnosed with prostate cancer (PCa) within their lifetime.^1^ Aggressive tumours progress to metastatic castration-resistant prostate cancer (mCRPC) which is responsible for the majority of PCa-related deaths.^2^ Other patients, however, will have clinically insignificant PCa, where the longevity and quality of a patient’s life is not adversely affected by PCa presence.^3^ Successfully determining between aggressive and clinically insignificant PCa is crucial to affording patients appropriate treatment. Current clinical diagnosis for PCa relies on multi-parametric MRI, PSA testing and transrectal ultrasound-guided biopsy (TRUS). ^4^ PSA screening in the UK is not currently implemented based on the limited benefits at diagnosing PCa on account of false negatives and false positives.^5^ Current testing for PCa is very limited prognostically and often leads to overtreatment of patients with clinically insignificant PCa. Another urgent biomarker requirement is for the accurate and early detection of resistance to hormonal therapies i.e. the development of castration resistance. This would facilitate the prompt discontinuation of ineffective therapies (with their significant sideeffects) and potentially adoption of new approaches.

Recent research has indicated that detection of circulating biomarkers including cell-free DNA, microRNAs, mRNAs and circulating tumour cells, present a minimally invasive alternative to current testing methods. ^6–9^ However, mRNA detection in particular, is often compounded by the limited specificity of qPCR assays.^10^ In addition, the relative low abundance in circulating biofluids of mRNA and its inherent lability can make this species a challenging yet potentially valuable dynamic biomarker for PCa prognosis. Detection of mRNA biomarkers at point-of-care (PoC) could provide rapid *in situ* responses to direct treatment options for PCa patients. Previous work has established several mRNAs of interest for PCa prognostics, including both androgen receptor (AR) variant 7 (AR-V7) and Yes-associated protein 1 (YAP1) mRNA.^11,12^ AR-V7 is deficient of the ligand binding domain (LBD) which normally makes the AR a ligand-activated transcription factor, as a result it is constitutively active. As such, AR-V7 presence in PCa patients is often associated with resistance to androgen deprivation therapy (ADT), the gold standard treatment for disseminated disease which targets the AR LBD.^13^ Across data extracted from twelve clinical trials, the proportion of mCRPC patients with detectable circulating AR-V7 mRNA is 18.3 %. ^14^ Detection of circulating AR-V7 mRNA in mCRPC patients treated with ADT, corresponded to reduced overall survival and progression free survival in these patients, supporting AR-V7 as clinically actionable mRNA for detection in the blood.^11^ YAP1, on the other hand, is commonly associated with the epithelial to mesenchymal transition in several types of cancers.^15–18^ In multiple PCa cell lines, YAP1 knockdown is associated with reduction in cellular motility, invasion and progression to metastatic phenotypes. ^19–21^ However, the *YAP1* gene is downregulated by late stage PCa-associated miR, miR-375-3p in mCRPC samples.^12,18^ Therefore, YAP1 poten-tially presents a temporal biomarker for progression from locally advanced PCa to mCRPC. Since the miR-375-3p - YAP1 pathway is implicated in docetaxel resistance, it could also direct treatment for mCRPC patients.^12^

qPCR is commonly referred to as the ‘gold standard’ for nucleic acid amplification tests, on account of its high accuracy and sensitivity. However, thermal cycling equipment crucial to qPCR experimentation is expensive and limited to use in specialized laboratories.^22^ As a result, qPCR experimentation is further compounded by transfer times to a laboratory. Alternative solutions for amplification tests are therefore required for point-of-care (PoC) prognostic and diagnostic tests. Loop-mediated isothermal amplification (LAMP), developed and optimised by Notomi *et al* and Nagamine *et al* respectively, is a rapid (<30 min) and sensitive DNA amplification technique.^23,24^ LAMP utilises six primers targeting eight specific DNA regions for exponential and isothermal amplification resulting in a high-yielding DNA assay.^23^ Reverse transcriptase LAMP (RT-LAMP) allows application of the technique to mRNA and has previously been used to detect mRNA in various diseases, including distinguishing dengue serotypes, prostate cancer antigen 3 for PCa diagnosis and more recently the N gene for SARS-CoV-2 virus detection. ^25–27^ Integration of LAMP assays with ion-sensitive field effect transistors (ISFETs) and unmodified complementary metal oxide semiconductor (CMOS) technology for Lab-on-Chip (LoC) detection of biomarkers has previously been successful. ^27–31^ RT-LAMP can be adjusted to result in a pH readout (RT-pHLAMP) during amplification events (i.e. a positive signal), which allows for compatibility with the pH-sensing ISFET for use in a microfluidic PoC device. ^32,33^ Double stranded DNA synthesis, which occurs in the RT-pHLAMP amplification event, releases a proton per nucleotide addition to the DNA strand.^34,35^

This work presents a method with bespoke primer selection and optimisation for the *de novo* development of RT-LAMP assays for the detection of AR-V7 and YAP1 mRNA. Adaptation of this assay for ISFET compatibility resulted in an accurate, sensitive (3×10^1^ copies per reaction) and rapid (<15 min) test for YAP1 and AR-V7 synthetic RNA presence. The assays were successfully tested on the ISFET LoC device presenting use of this device for PoC. Validation of this assay and the LoC device was confirmed with detection of AR-V7 and YAP1 mRNA extracted from PCa cell lines. The development of this biosensor and these assays present the potential for PoC prognostics, where clinicians can rapidly adjust treatment options for PCa patients.

## Results

### RT-qLAMP and RT-pHLAMP assay optimisation for AR-V7 and YAP1 detection

Initial optimisation of the RT-qLAMP assay rendered the primers shown in Table 1. Different lengths of the front inner primer and back inner primer were tested to ensure optimal time to positive (TTP) values were achieved. The AR-V7 primers specifically targeted a region in cryptic exon 3 to avoid amplification of the fulllength androgen receptor (AR-FL) mRNA.^36^ Since mRNAs present in the blood are often fragmented, synthetic RNA fragments of both AR-V7 and YAP1 target regions (374 bp and 355 bp lengths respectively) were synthesized for initial assay development.^37,38^ Both the AR-V7 and YAP1 RT-qLAMP assays achieved linear detection of 3×10^7^ to 3×10^2^ copies of synthetic RNA per reaction in under 17 min (Figure 2).

**Table 1:**
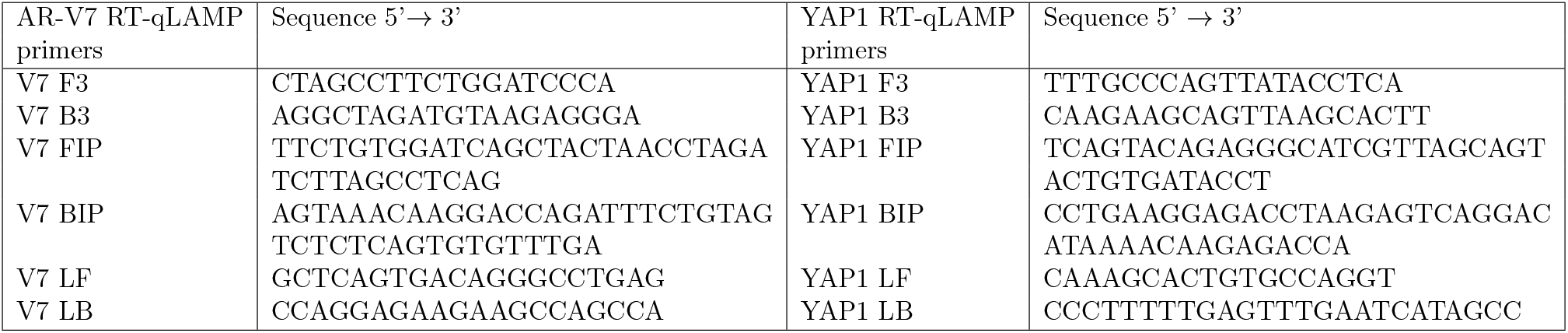
Primer sequences for both AR-V7 and YAP1 RT-qLAMP and RT-pHLAMP assays.

**Figure 1:**
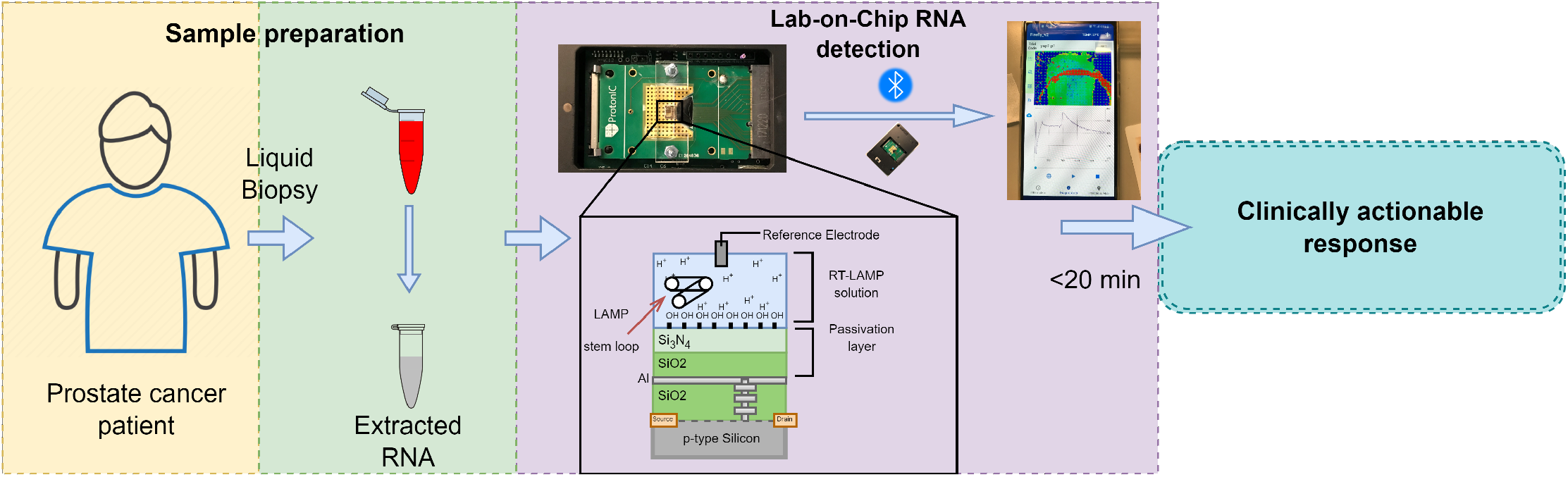
Prospective workflow from liquid biopsy extraction from a PCa patient to a clinically actionable response via mRNA detection using an ISFET biosensor and an optimised RT-LAMP assay.

**Figure 2:**
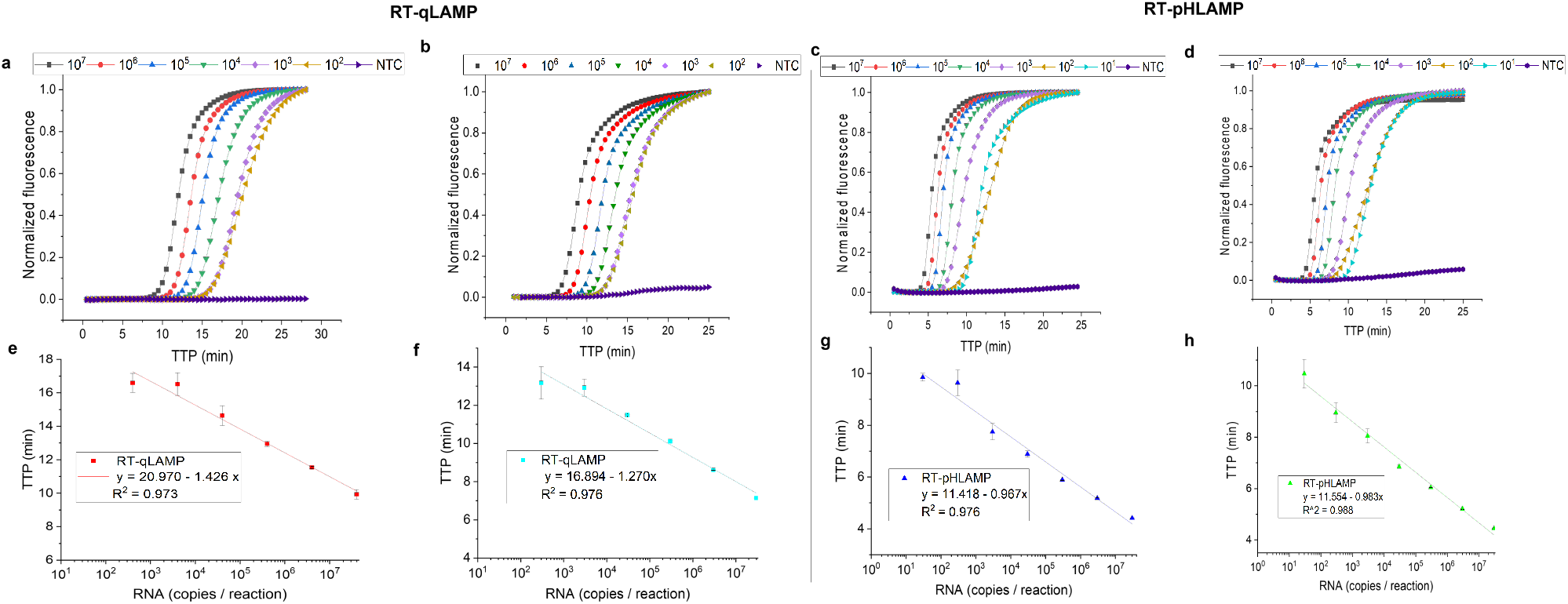
**(a-d)** Sigmoidal amplification curves of RT-qLAMP and RT-pHLAMP assays detecting AR-V7 and YAP1 synthetic RNA. Synthetic RNA concentrations varied from 3×107 to 3×102 copies per reaction for each RT-qLAMP reaction and 3×107 to 3×101 copies per reaction for each RT-pHLAMP reaction. **(a)** Amplification curve of the RT-qLAMP assay detecting synthetic AR-V7 RNA. **(b)** Amplification curve of the RT-qLAMP assay detecting synthetic YAP1 RNA. **(c)** Amplification curve of the RT-pHLAMP assay detecting synthetic AR-V7 RNA. **(d)** Amplification curve of the RT-pHLAMP assay detecting synthetic YAP1 RNA. **(e-h)** Standard curves of RT-qLAMP and RT-pHLAMP detection of synthetic AR-V7 and YAP1 RNA at varying concentrations. These graphs include linear regressions, the coefficient of determinations of each assay and error bars displaying one standard deviation. **(e)** Standard curve of the RT-qLAMP assay detecting synthetic AR-V7 RNA. **(f)** Standard curve of the RT-qLAMP assay detecting synthetic YAP1 RNA. Data averaged across two experiments. **(g)** Standard curve of the RT-pHLAMP assay detecting synthetic AR-V7 RNA. **(h**) Standard curve of the RT-pHLAMP assay detecting synthetic YAP1 RNA. Data averaged across two experiments.

In order to generate a pH readout for IS-FET compatibility, the RT-qLAMP assays were adjusted as previously described to omit tris(hydroxymethyl)-aminomethane (tris), the pH buffering agent present in Isothermal Amplification Buffer (New England Biolabs).^32^ Betaine was further omitted in the augmented assay to equate for lyophilisation compatibility. The resulting RT-pHLAMP assays subsequently showed a sensitivity of 3×10^1^ RNA copies per each reaction (Figure 2). The standard curves of these reactions presented coefficients of determination (R^2^) of 0.976 and 0.988 for the AR-V7 and YAP1 RT-pHLAMP assays respectively, which indicates the potential of these assays for accurate quantification of RNA per sample. TTP values for the pH sensitive reactions were significantly reduced: the average TTP for detection of 3×10^2^ copies of synthetic AR-V7 RNA was 16.6 min in RT-qLAMP and 9.6 min in RT-pHLAMP. This is likely due to the increased optimisation of the RT-pHLAMP assay, allowing for faster TTP values. Detection from 3×10^7^ to 3×10^1^ copies of RNA was achieved in under 12 min for both RT-pHLAMP assays.

Specificity of the AR-V7 RT-pHLAMP reaction was confirmed by spiking the assays with a synthetic RNA fragment present in the AR-FL LBD (Supplementary Figure 4). Primers detecting this AR-FL region were developed to confirm its the presence in these spiked assays (Supplementary Figure 5). No amplification occurred between the AR-FL synthetic RNA and AR-V7 primers after the reaction was terminated at 35 min. A serial dilution experiment for AR-V7 detection spiked with AR-FL then took place. These results indicate that amplification of the AR-V7 RT-pHLAMP assay only occurred with the presence of AR-V7 mRNA. In this instance, the sensitivity of the reaction was reduced to 3×10^2^ copies, indicating that presence of off-target RNA decreased the efficiency of the RT-pHLAMP assay.

### Validation of AR-V7 and YAP1 RT-pHLAMP specificity with extracted RNA from prostate cancer cell lines

Extracted RNA from PCa cell lines 22Rv1 and DU145 was utilised to confirm the detection of endogenous YAP1 and AR-V7 mRNA. 22Rv1s have previously been reported as AR-V7 mRNA positive whilst DU145s show little to no AR-V7 expression.^39^ Five individual 22Rv1 RNA samples rendered an average TTP of 8.17 *±* 0.54 min with 1 ng of RNA per reaction (Figure 3 **b**). In contrast, 1 ng per reaction of extracted RNA from DU145s rendered no fluorescent signal after 35 min, indicating no amplification had taken place. These findings suggest the AR-V7 RT-pHLAMP assay is specific to AR-V7 mRNA in patient-derived cell lines.

**Figure 3:**
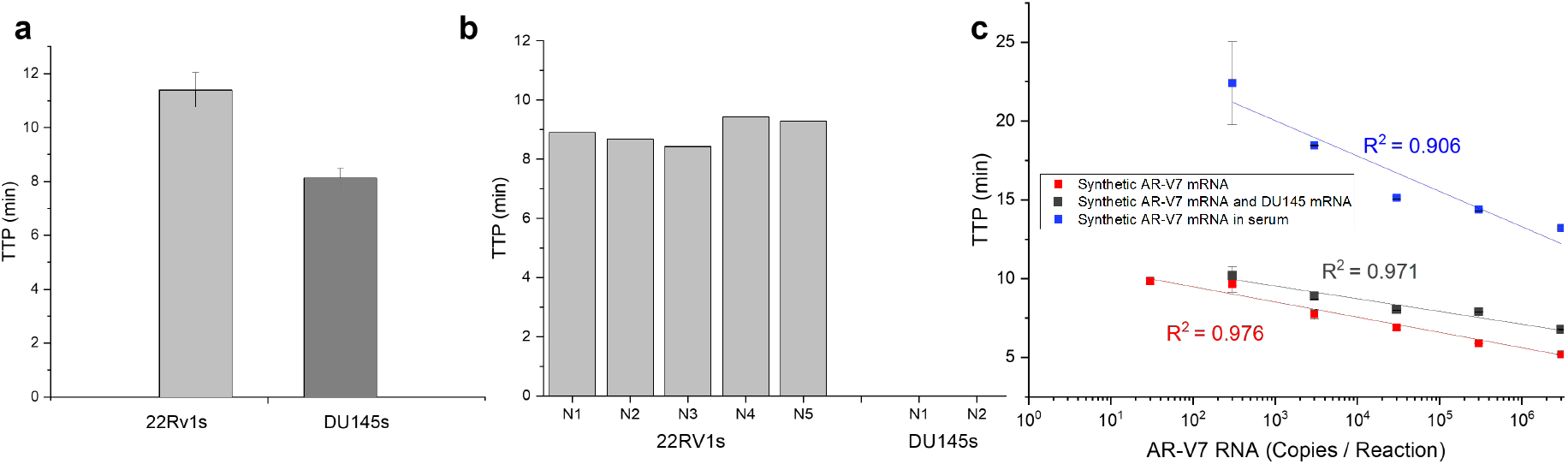
**(a)**: The variation in TTP in the YAP1 RT-pHLAMP assay with 1 ng of extracted mRNA from two PCa cell lines, DU145s and 22Rv1s. **(b)**: The variation in TTP in the AR-V7 RT-pHLAMP assay with 1 ng of extracted RNA from AR-V7 positive cell line, 22Rv1 and the AR-V7 negative cell line, DU145. **(c)**: The standard curves for multiple AR-V7 RT-pHLAMP experiments including the unmodified synthetic AR-V7 mRNA assay, the assay spiked with off-target DU145 mRNA and the assay containing mixed male serum.

In order to determine if off-target RNA affected the efficiency of the RT-pHLAMP assay, synthetic AR-V7 mRNA was spiked with 1 ng of DU145 mRNA. A serial dilution experiment was then conducted (Figure 3 **c**). These results suggest that the presence of off-target RNA marginally increased the TTP values at various concentrations in the AR-V7 RT-pHLAMP assay. In addition, reliable and quantitative detection of synthetic RNA was achieved down to 3×10^2^ copies per reaction. Utilising the standard curve generated from this experiment, the average copy number of AR-V7 mRNA per 1 ng of RNA is 1.6 × 10^5^ copies.

To indicate if both the RT-pHLAMP assays for AR-V7 and YAP1 mRNA were feasible for detection of circulating mRNA in the blood, assays including mixed male serum were conducted (Figure 3 **c** and Supplementary Figure 6 respectively). The limit of detection in these experiments was also 3×10^2^ copies per reaction although TTP values were increased. pH values for these reactions indicated that no pH change took place, likely due to the carbonic acid / bicarbonate buffer system present in the blood.^40^ Integration of serum samples directly on to the LoC platform would subsequently require further optimisation, outside of the scope of this study.

YAP1 mRNA presence was also tested in RNA extracted from 22Rv1 and DU145 cell lines. High expression of YAP1 mRNA concentration has previously been recorded in DU145 cells.^19^ The RT-pHLAMP assay detected YAP1 mRNA presence in 8.08 *±* 0.41 min at 1 ng per reaction across RNA extracted from two DU145 cell line samples. YAP1 presence was additionally detected in 22Rv1 RNA samples, at an increased TTP of 11.7 *±* 0.68 min. miR-375 is highly expressed in 22Rv1 cell lines and targets YAP1 mRNA, resulting in its downregulation.^18^ As such, the variation in TTP values for 22Rv1 and DU145 extracted RNA samples corresponds to the expected concentrations of YAP1 mRNA in these cell lines. A Welch’s unequal variances *t* -test was used to confirm the significance of this data (t = 14.47, *p* < .001). RT-qPCR assays (Supplementary Figure 8) confirmed the high concentration of YAP1 mRNAs in DU145s and lower concentration in 22Rv1s (t = 8.15, *p* < .001). Also as expected, no amplification curves were seen in DU145s with the AR-V7 RT-qPCR assay, while fast amplification was observed in the 22Rv1 cell line (Supplementary Figure 7). This indicates that the RT-pHLAMP assay data corresponds well with the gold standard of nucleic acid amplification tests.

### Implementation of AR-V7 and YAP1 RT-pHLAMP assays onto the Lab-on-Chip platform

The developed RT-pHLAMP assays were subsequently imtegrated into the Lab-on-Chip which utilised ISFET sensors to detect the rate of pH change. Double-stranded DNA synthesis, which occurs during the RT-LAMP amplification event (in positive samples), releases a proton per each nucleotide addition.^34,35^ The subsequent change in pH of the unbuffered RT-pHLAMP solution is detected by the ISFET and recorded by a mobile phone.

Synthetic YAP1 and AR-V7 RNA samples were successfully detected at a concentration of 3×10^6^ copies per reaction. TTP values were slightly increased on the LoC platform, likely due to non-optimal conditions for the RT-pHLAMP assay in the acrylic reaction chamber. These increased values are still significantly reduced (indicating more rapid detection) relative to the PCR gold standard for nucleic acid amplification tests. The averaged TTP value across triplicate experiments for detection of YAP1 and AR-V7 synthetic RNA at 3×10^6^ copies per reaction was 7.25 *±* 0.62 min and 7.11 *±* 0.65 min respectively. Figure 4 shows the implementation of the AR-V7 RT-pHLAMP assay onto the microchip. Postprocessing of the voltage readout is required to subtract the inherent drift present in ISFET biosensors. ^41^ The voltage output is converted to proton count and sigmoidal fitting is then carried out to return the amplification curve illustrated in Figure 4.

**Figure 4:**
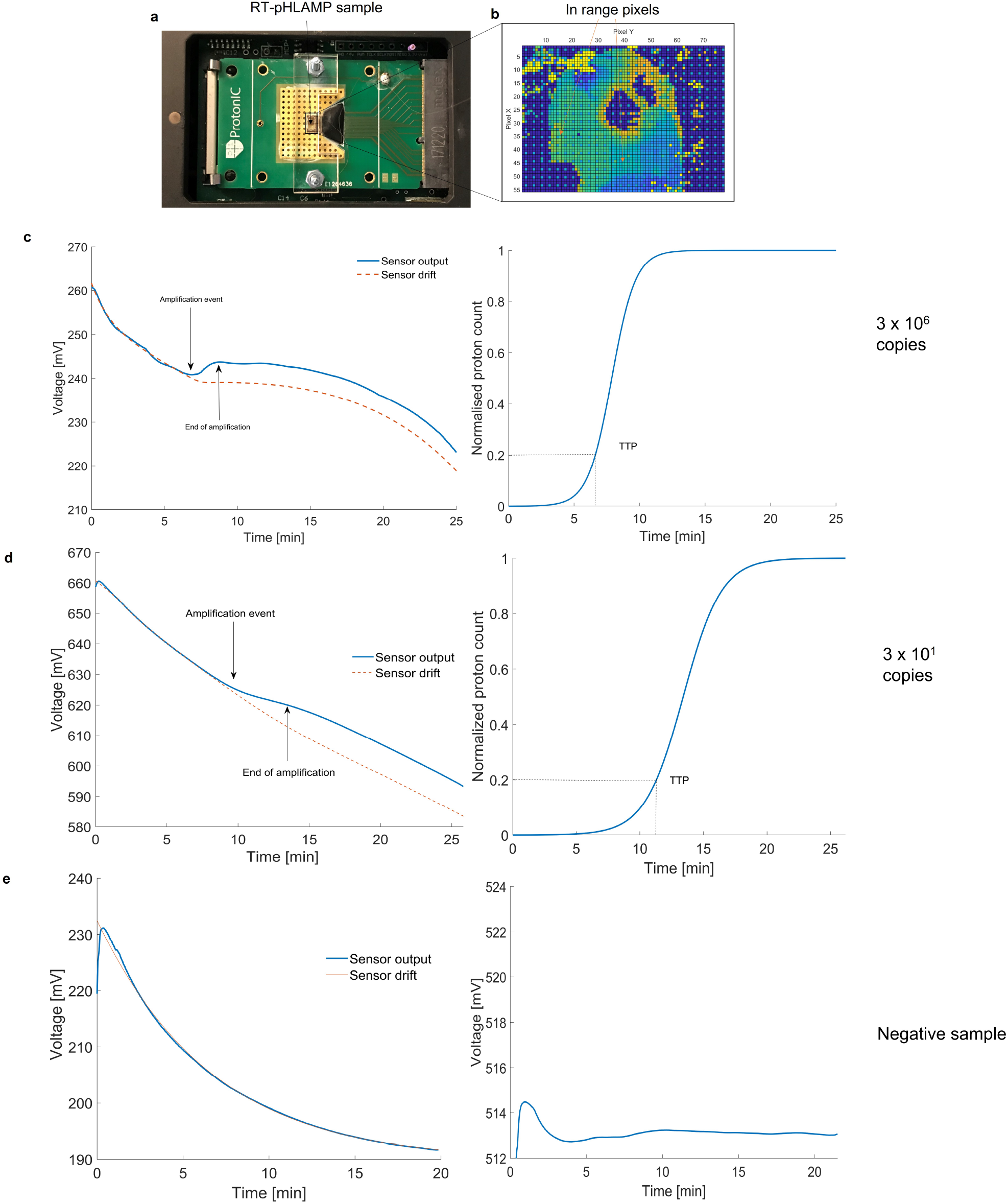
Illustration of RT-pHLAMP implementation onto the LoC platform. **(a)**: ISFET microchip setup with an acrylic manifold and RT-pHLAMP sample loaded onto the microchip. **(b)**: The array of the ISFET microchip once the experiment was initiated. In range pixels are shown in green / light blue. Dark blue and red indicate pixels that are out of range for pH detection. **(c)**: The ISFET sensor output graph (left) and sigmoidal-fitted amplification curve (right) of a positive AR-v7 sample on the ISFET microchip (3 × 106 copies per reaction). **(d)**: Detection of 3 × 101 copies of synthetic AR-v7 RNA with the ISFET biosensor. The ISFET biosensor output graph (left) and the amplification curve with sigmoidal fitting (right) are shown here. **(e)**: The ISFET sensor output graph (left) and amplification curve (right) of a negative AR-v7 sample on the ISFET microchip. No sigmoidal fitting was performed for this experiment on account of the negative signal.

Once the detection of 3×10^6^ copies of both AR-V7 and YAP1 synthetic RNA had taken place with the LoC device, the limit of detection was tested at 3×10^1^ copies. For both of the RT-pHLAMP assays the pH change at 3×10^6^ and 3×10^1^ copies were similar, likely due to the DNA production being the same at both concentrations. Figure 4 additionally shows the amplification of 3×10^1^ copies of AR-V7 synthetic mRNA on the LoC device. Here, the TTP values were 10.88 *±* 0.95 min for the AR-V7 RT-pHLAMP assay and 11.50 *±* 0.98 for the YAP1 RT-pHLAMP assay. This illustrates that the sensitivity of the LoC device is comparable to the benchtop RT-pHLAMP assays.

### Detection of YAP1 and AR-V7 mRNA from patient-derived cell lines on the Lab-on-chip platform

Once it had been determined that these assays were compatible with the ISFET biosensor, detection of AR-V7 and YAP1 mRNA present in RNA extracted from 22Rv1 and DU145 cells was assessed. As confirmed in the previous benchtop RT-pHLAMP and RT-qPCR assays, 22Rv1 cells are AR-V7 positive and DU145s contain high levels of YAP1. Figure 5 shows the ISFET detection of AR-V7 and YAP1 in the two cell lines. Here, no positive signal is detected for AR-V7 in the DU145 cell line, mirroring the expression shown in the RT-qPCR-based assay and the relevant literature.^39^ Contrastingly, detection of 1 ng of AR-V7 mRNA per reaction was achieved in 8.48 *±* 1.43 min in extracted RNA from 22Rv1s on the LoC device. Figure 5 **(a)** illustrates the comparison between the LoC device and the benchtop assays for detection of AR-V7 and YAP1 mRNA. The LoC values are largely comparable to the pH change and TTP values of the benchtop assay, indicating that the LoC device is a robust method for AR-V7 and YAP1 detection in PCa cell lines. YAP1 mRNA detection occurs in 8.01 *±*0.64 min with DU145 mRNA and 13.22 *±* 1.59 min with 22RV1 mRNA. The change in TTP values between the two PCa cell lines on average is 5.22 min, which is increased relative to the benchtop assay. As such, it provides a greater distinction between YAP1 mRNA concentrations within 22Rv1 and DU145 cell lines.

**Figure 5:**
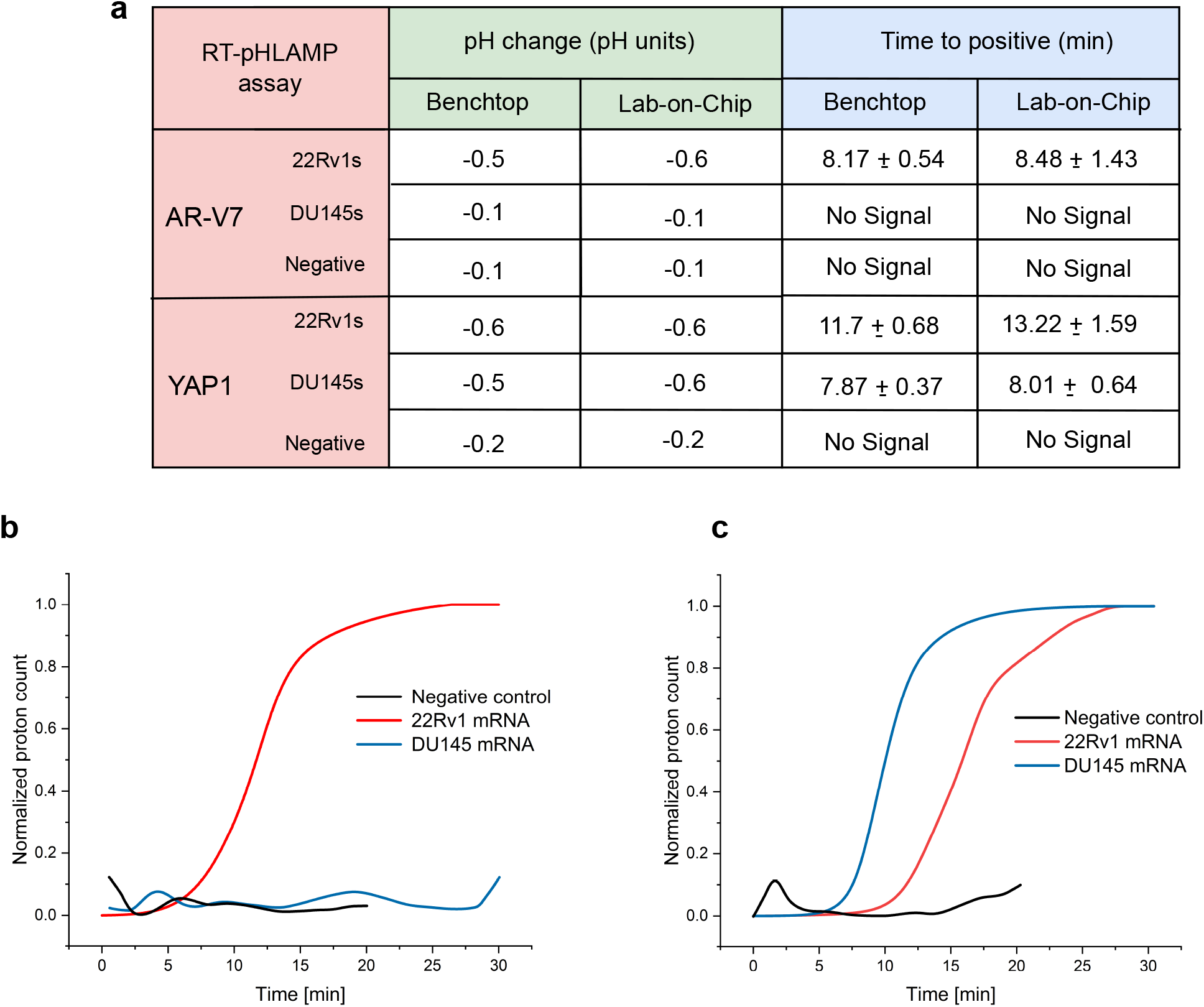
**(a)**: TTP and pH change values of the benchtop and LoC RT-pHLAMP assays. Benchtop assays were terminated after 35 min, LoC positives were terminated after 30 min and LoC negatives after 20 min. **(b)**: The sigmoidal-fitted averages of AR-V7 mRNA in 22Rv1s, DU145s and negative samples using the RT-pHLAMP assay. Graphs are the average values of triplicate assays. **(c)**: The sigmoidal-fitted averages of YAP1 mRNA in 22Rv1s, DU145s and negative samples using the RT-pHLAMP assay. Graphs represent the average values of triplicate assays.

**Figure 6:**
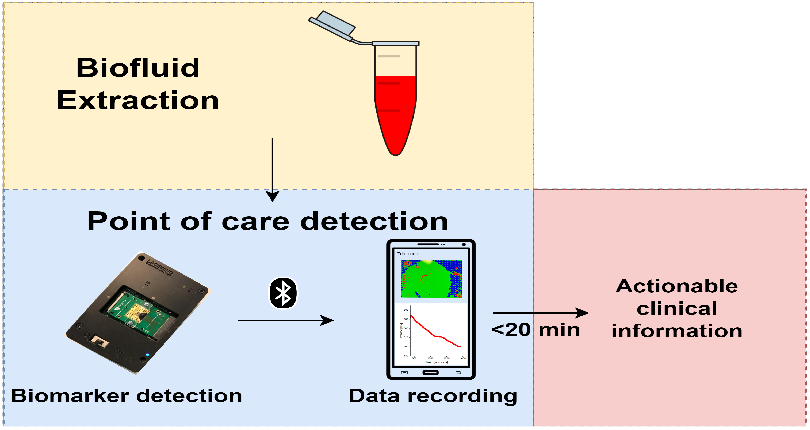
ToC graphic with dimensions 7 cm x 3.5 cm. i.e. below the constraints of 9 cm x 3.5 cm.

## DISCUSSION

This paper presents a foundation for the LoC detection of circulating mRNA in PCa. The two novel assays judiciously developed for this work are, to the authors’ knowledge, the first RT-qLAMP experiments for the detection of AR-V7 and YAP1 mRNA. Authentication of AR-V7 and YAP1 detection was confirmed with extracted RNA from PCa cell lines and RT-qPCR. These RT-pHLAMP assays produced a suitable pH change for use with CMOS technology containing an array of ISFET sensors. This compatibility resulted in a LoC device with potential for direct PoC usage. Detection of synthetic RNA was achieved at a sensitivity of 3 × 10^1^ copies per reaction for both markers. The RT-pHLAMP reactions, on account of their isothermal nature, remove the necessity of specialised and expensive thermal cycling equipment required for RT-qPCR experiments. Further development of these assays for the detection of circulating mRNA directly in serum would further increase their potential for rapid prognostics.

The PROPHECY study, completed in 2019, has confirmed that the presence of circulating AR-V7 mRNA associates with a lower progression-free survival and overall survival in mCRPC patients treated with enzalutamide and abiraterone.^11^ This illustrates that the presence of circulating AR-V7 mRNA could be used to monitor mCRPC patients and direct treatment options. Therefore, PoC detection of AR-V7 through this novel assay could show clinical benefit to mCRPC patients.

YAP1 concentration can be distinguished in the quantitative RT-pHLAMP assay and detected using the LoC device with varying TTP values in two PCa cell lines. High YAP1 concentration can be illustrative of PCa tumours progressing to EMT while low YAP1 concentration could indicate advancement of mCRPC towards docetaxel resistance. In conjunction, AR-V7 and YAP1 mRNA detection on a LoC device, could result in clinically actionable information, obtainable rapidly (<20 min), sensitively and directly in the clinic. Further evaluation utilising blood samples from PCa patients will be required to confirm the validity of these assays for use directly in hospitals. Progression in sample preparation allowing for direct detection of circulating markers in the blood in RT-pHLAMP reactions will expedite the time taken from biofluid extraction to a prognosis using this PoC device.

Further detection of a larger range of circulating nucleic acid biomarkers could create a multiplex LoC device to serve as a prognostic test to personalize medication for PCa patients. The development of more RT-pHLAMP assays in conjunction with the ISFET LoC device, could result in a robust handheld device for rapid, reliable and simultaneous detection of multiple circulating prognostic PCa biomarkers.

## Materials and methods

### Synthesis of synthetic RNA targets

RNA fragments of AR-V7 and YAP1 sequences were synthesized from DNA gblocks (Integrated DNA Technologies) utilising the HiScribe™ T7 Quick High Yield RNA Synthesis Kit (NEB) according to the manufacturer’s instructions including the DNase step. Stock concentrations were maintained at 3×10^10^ copies per *µ*L and stored at -80 °C in preparation for experiments.

### RT-qLAMP experiments

All reactions were completed in triplicate. Each 10 *µ*L experiment contained: 1 *µ*L 10x isothermal buffer (New England Biolabs (NEB)), 0,6 *µ*L MgSO_4_ (100 mM stock), 1.4 *µ*L dNTPs (10 mM stock of each nucleotide), 0.6 *µ*L BSA (20 mg / mL stock), 0.8 *µ*L betaine (5 M stock), 0.25 *µ*L SYTO 9 green (20 *µ*L stock), 0.25 *µ*L NaOH (0.2 M stock), 0.042 *µ*L Bst 2.0 DNA polymerase (120,000 U / ml stock, NEB), 0.1 Ribolock RNAse Inhibitor *µ*L (40 U / *µ*L stock, ThermoFisher), 0.3 *µ*L Warmstart® RTx reverse transcriptase (15,000 U / ml, NEB), 1 *µ*L 10x LAMP primer mix (20 *µ*M FIP and BIP, 10 *µ*M LB and LF, 2.5 *µ*M F3 and B3), 1 *µ*L RNA sample and the remaining solution was topped up to 10 *µ*L with nuclease-free water. Reactions were conducted at 63°C for 35 min. One melting curve from 63 °C to 97 °C was conducted to confirm the specific amplification of the reaction at a ramp of 0.2 °C / s. Reactions were conducted with a Lightcycler® 96 instrument (Roche Diagnostics) in 96 well plates.

### RT-pHLAMP experiments

All reactions were completed in triplicate. Each 10 *µ*L experiment contained: 1 *µ*L customized isothermal buffer, 0.5 *µ*L MgSO_4_ (100 mM stock), 1.4 *µ*L dNTPs (10 mM stock of each nucleotide), 0.6 *µ*L BSA (20 mg / ml stock), 0.25 *µ*L SYTO 9 green (20 *µ*M stock), 0.25 *µ*L NaOH (0.2 M stock), 0.042 *µ*L Bst 2.0 Warmstart® DNA polymerase (120,000 U / mL stock, NEB), 0.3 *µ*L Warmstart ® RTx reverse transcriptase (15,000 U / ml stock, NEB), 1 *µ*L 10x LAMP primer mix (20 *µ*M FIP and BIP, 10 *µ*M LB and LF, 2.5 *µ*M F3 and B3), 1 *µ*L RNA sample and the remaining solution was topped up to 10 *µ*L with nuclease-free water. For serum experiments, the RNA sample was diluted in mixed male serum (Sigma Aldrich) and 1 *µ*L of that solution was added to the reaction. Reactions were conducted at 63°C for 35 min. One melting curve from 63 °C to 97 °C was conducted to confirm the specific amplification of the reaction at a ramp of 0.2 °C / s. Reactions were conducted with a Lightcycler ® 96 instrument (Roche Diagnostics) in 96 well plates. Reactions were scaled up to either 12 *µ*L or 20 *µ*L reactions for implementation onto the LoC device, proportions of each reagent were kept the same.

### RT-qPCR experiments

All reactions were completed in triplicate. RT-qPCR reactions were completed in two steps. 50 ng mRNA samples were initially converted to cDNA with a RevertAid First Strand cDNA synthesis kit (ThermoFisher Scientific) as per the manufacturer’s instructions including the optional step for GC rich regions. cDNA was used immediately for qPCR assays. qPCR experiments were conducted in 10 *µ*L quantities and contained the following: 5 *µ*L Fast SYBR® Green Master Mix (Applied Biosystems), 2 *µ*L cDNA sample, 0.5 *µ*L forward primer (250nM, 5 *µ*M stock), 0.5 *µ*L reverse primer (250nM, 5 *µ*M stock), Nuclease-free water was added to make the reaction volume up to 10 *µ*L. Reactions were aliquoted into a 96 well plate for analysis with a StepOnePlus™Real-Time PCR system (Applied Biosystems). Reactions were initially heated to 95 °C for 20 s. The cycling stage including heating at 95 °C for 3 s followed by 60 °C for 30 s. The cycling stage was repeated for 40 cycles. Melting curves were conducted with heating to 95 °C for 15 s followed by 60 °C for 1 min.

### Translation of RT-pHLAMP onto Lab-on-Chip device

The LoC system detects changes in proton concentration on the interface of the RT-pHLAMP assay solution with the passivation layer (Si_3_N_4_). The ISFET array is comprised of 56 × 78 ISFET pixels (4368 individual sensors, 2 × 4 *mm*).42 Temperature was maintained at 63 °C with a peltier heating module contacting the underside of the cartridge. The LoC device was battery-powered and data was sent to an Android phone through a bluetooth connection. Data extracted from the mobile phone was run through a MATLAB (R2021b) algorithm designed to spot for amplification events. The RT-pHLAMP assay solutions were housed in an acrylic manifold with either 12 *µ*L or 20 *µ*L sized chambers. Adhesive gaskets comprised of Tesa® double-sided smooth lamination filmic tape sealed the acrylic manifold to the cartridge. A 0.03 mm chlorodised silver wire served as a Ag/AgCl reference electrode. This electrode was in contact with the assay solution and was placed between the adhesive gasket and the microchip’s surface. Nucleasefree Water was added to the chamber of the manifold for the first 700 s to equilibrate the system and set a common voltage across the ISFET array. The water was then extracted and the RT-pHLAMP reaction mixture was added. All samples that contained synthetic RNA or extracted RNA were run for 30 min after the addition of the RT-pHLAMP reaction. Negative controls contained nuclease-free water instead of RNA and were run for 20 min after the addition of the RT-pHLAMP reaction. All reactions were completed in triplicate. Confirmation of amplification presence in these reactions was confirmed with a Qubit 3.0 fluorometer (Invitrogen). Measured pH values postreaction were conducted with a microFET pH probe (Sentron).

### RNA extraction from prostate cancer cell lines

22Rv1 and DU145 cell lines were cultured in T75 flasks with RPMI-1640 containing FBS (10%) and L-Glutamine (5 mM). Cells were passaged at 70 % confluency to maintain optimal growth and kept to under 10 passages post-thawing. For RNA extraction, cells were harvested at 70 % confluency and spun down to remove media. For the 22Rv1 cells, RNA extraction was performed using the Total RNA Miniprep Kit (Monarch) as per manufacturer’s instructions including the DNase I digestion step. For DU145 cells, RNA extraction was performed using the RNeasy Mini Extraction kit (Qiagen) as per the manufacturer’s instructions. In both cases, RNA was eluted in 50 *µ*L and RNA quantity/quality measured using a Nanodrop D1000. Extracted RNA was stored at -80 °C until use.

### Statistical analyses

The Welch’s *t* test was chosen to determine the statistical significance of YAP1 RT-pHLAMP TTP values and YAP1 qPCR Cq values in the 22Rv1 and DU145 cell lines experiments. This test is commonly utilised in scenarios where the two compared datasets have different variance or different sample sizes.43 The null hypothesis in each case was that the mean TTP or Cq value was the same between the 22Rv1 and DU145 prostate cancer cell lines.

The calculation for degrees of freedom (*v*) for Welch’s *t* -test is shown below (Equation **(1)**), where *s*_1_ and *s*_2_ are the standard deviations of the two datasets and *N*_1_ and *N*_2_ are the number of samples per dataset.

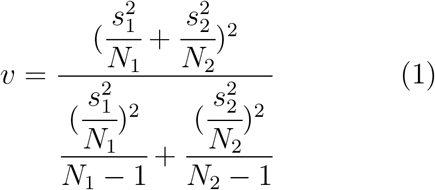

The equation for *t* value for the Welch’s *t* -test of unequal variance is shown below (Equation **(2)**), where 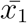 and 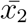are the mean values of the two datasets.

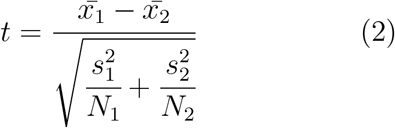

The null hypothesis was rejected when *p* < .05.

## Acknowledgement

The authors thank members of the Georgiou and Bevan Laboratories as well as Dr Sylvain Ladame for discussions around this work.

## Supporting Information Available

The following file is available free of charge.

- Supplementary file 1: Includes further experimental data referenced in the main manuscript.

