## Supplementary information and data for "Detection of YAP1 and AR-V7 mRNA for Prostate Cancer prognosis using an ISFET Lab-On-Chip platform"

**4 AR-V7 specificity** **7**

**5 YAP1 RT-pHLAMP and serum reaction** **9**

**6 qPCR data for AR-V7 mRNA in PCa cell lines** **10**

**7 qPCR data for YAP1 mRNA in PCa cell lines** **11**

**8 qPCR primer sequences** **11**

**9 Cq value determination** **12**

**10 Materials and Methods** **12**

### 1 DNA gblocks

Each gblock (Integrated DNA Technologies) contains the 5'-TAATACGACTCACTATAGG-3' promoter region for T7 polymerase. These gblocks were transcribed into RNA for use in synthetic RNA experiments as described under the Materials and Methods section.

#### 1.1 AR-V7 gblock

Accession number: NM_001348061
5'-TAATACGACTCACTATAGGGGACTCAAGGTGTCACCTTGGACAAGAAG
CAACTGTGTCTGTCTGAGGTTCCTGTGGCCATCTTTATTTGTGTATTAGGC

AATTCGTATTTCCCCCTTAGGTTCTAGCCTTCTGGATCCCAGCCAGTGACC
TAGATCTTAGCCTCAGGCCCTGTCACTGAGCTGAAGGTAGTAGCTGATCCA
CAGAAGTTCAGTAAACAAGGACCAGATTTCTGCTTCTCCAGGAGAAGAAGC
CAGCCAACCCCTCTCTTCAAACACACTGAGAGACTACAGTCCGACTTTCCC
TCTTACATCTAGCCTTACTGTAGCCACACTCCTTGATTGCTCTCTCACATC
ACATGCTTCTCTTCATCAGTTGTAAGCCTCTCATT-3'

##### 1.2 YAP1 gblock

Accession number: NM_001130145
5'-TAATACGACTCACTATAGGGGTGCTGCCATTAAAGGCAGCTGTTCTAG
AGTTTCAGTCACCTAAGTACACCCACAAAACAATATGAATATGGAGATCTT
CCTTTACCCCTCAACTTTAATTTGCCCAGTTATACCTCAGTGTTGTAGCAG
TACTGTGATACCTGGCACAGTGCTTTGATCTTACGATGCCCTCTGTACTGA
CCTGAAGGAGACCTAAGAGTCCTTTCCCTTTTTGAGTTTGAATCATAGCCT
TGATGTGGTCTCTTGTTTTATGTCCTTGTTCCTAATGTAAAAGTGCT-
TAACTGCTTCTTGGTTGTA TTGGGTAGCATTGGGATAAGATTTTAACTGGG
TATT CTTGAATTGCTTTTAC-3'

##### 1.3 AR-FL gblock

Accession number: NM_000044
5'-TAATACGACTCACTATAGGCTCCGTGCAGCCTATTGCGAGAGAGCTGCA
TCAGTTCACTTTTGACCTGCTAATCAAGTCACACATGGTGAGCGTGGACTTT
CCGGAAATGATGGCAGAGATCATCTCTGTGCAAGTGCCCAAGATCCTTTCTG
GGAAAGTCAAGCCCATCTATTTCCACACCCAGTGAAGCATTGGAAACCCTAT
TTCCCCACCCCAGCTCATGCCCCCTTTCAGATGTCTTCTGCCTGTTATAACT
CTGCACTACTCCTCTGCAGTGCCTTGGGGAATTTCCTCTATTGATGTACAGT
CTGTCATGAACATGTTCCTGAATTCTATTTGCTGGGCTTTTTTTTTCTCTTT
CTCTCCTTTCTTTTTCTT-3'

#### 2 AR-FL primers

F3 - 5'-AGACTCTCTCCAGACAGC-3'
B3 - 5'-GACTTTAAGTTTTGGATTTGATCTG-3'
LF - 5'-TTCCAGGGCTATGCAGGGG-3'
LB - 5'-GTTTGACCCACTACAAGGGGT-3'
FIP - 5'-TTCGTAGACAGTCAGCCTCACTACCCGAGCATGGCCCC-3'
BIP - 5'- GCCAAGGGAGTGGTTTTTTCCTGATTCCCATGAC-3'

#### 3 Primer optimisation experiments

### 3.1 AR-V7

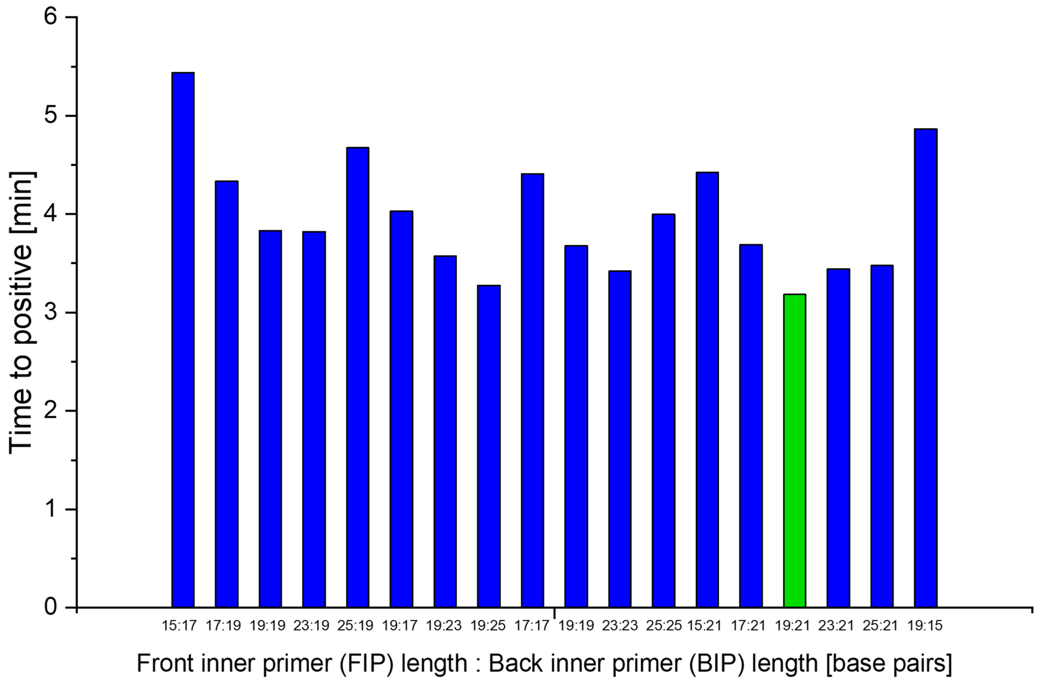

Figure 1: Optimisation RT-qLAMP experiment determining the fastest combination of FIP and BIP base pair values for the AR-V7 synthetic target. The fastest combination is shown in green and it's sequence is shown in the main manuscript. 3 x $10^8$ copies of synthetic RNA are detected in these reactions.

##### 3.2 YAP1

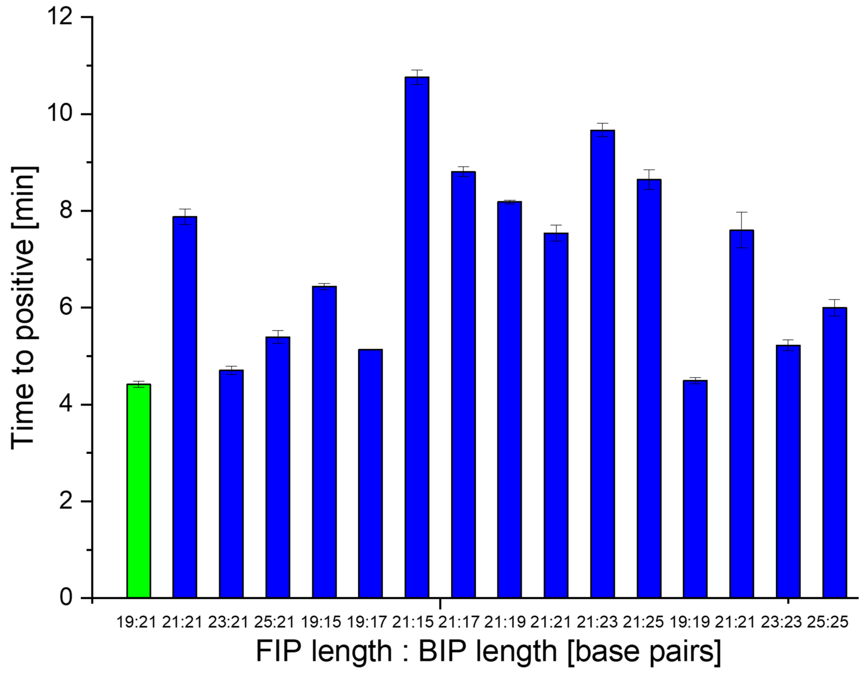

Figure 2: Optimisation RT-qLAMP experiment determining the fastest combination of FIP and BIP base pair values for the YAP1 synthetic target. The fastest combination is shown in green and it's sequence is shown in the main manuscript. 3 x $10^8$ copies of synthetic RNA are detected in these reactions.

### 3.3 AR-FL

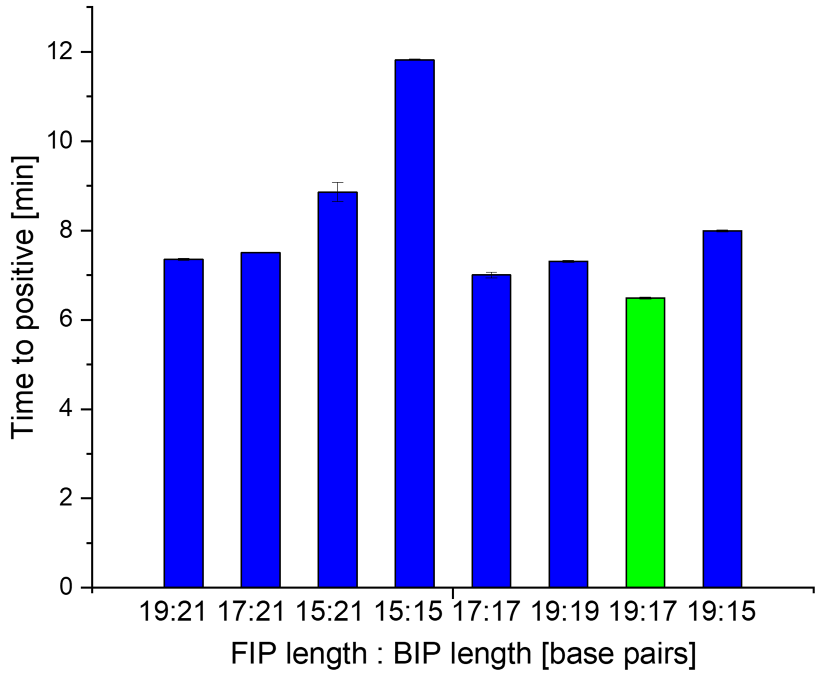

Figure 3: Optimisation RT-qLAMP experiment determining the fastest combination of FIP and BIP base pair values for the YAP1 synthetic target. The fastest combination is shown in green and it's sequence is shown above under AR-Fl primers. 1 x $10^8$ copies of synthetic RNA are detected in these reactions.

### 4 AR-V7 specificity

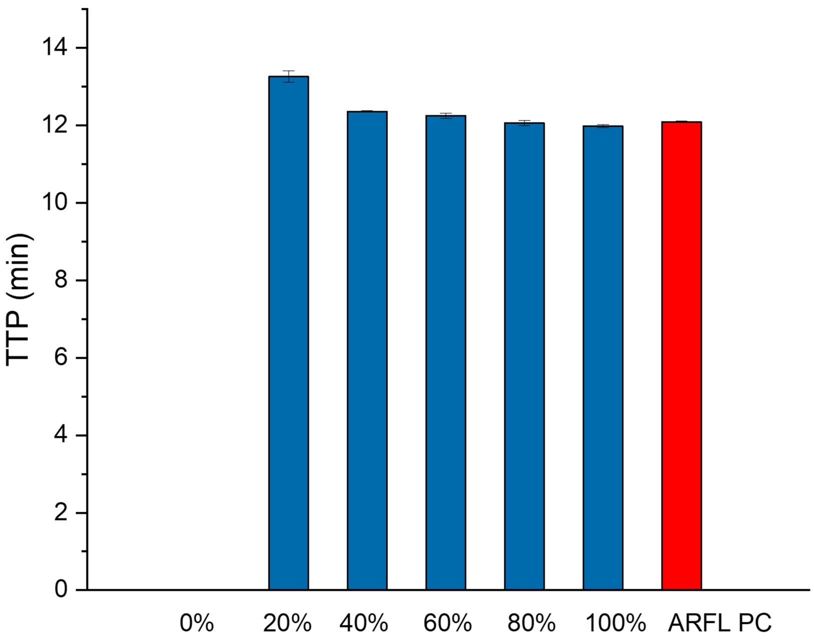

Figure 4: This graph shows the data of the AR-V7 RT-qLAMP assay spiked with AR-FL synthetic RNA. Percentages shown on the x axis indicate the relative quantity of AR-V7 synthetic RNA in the assay. The total RNA copies per reaction was 1 $x10^5$ copies. For example in the 20 % assay $2x10^4$ copies of AR-V7 synthetic RNA were present and $8x10^4$ copies of AR-FL synthetic RNA were present.

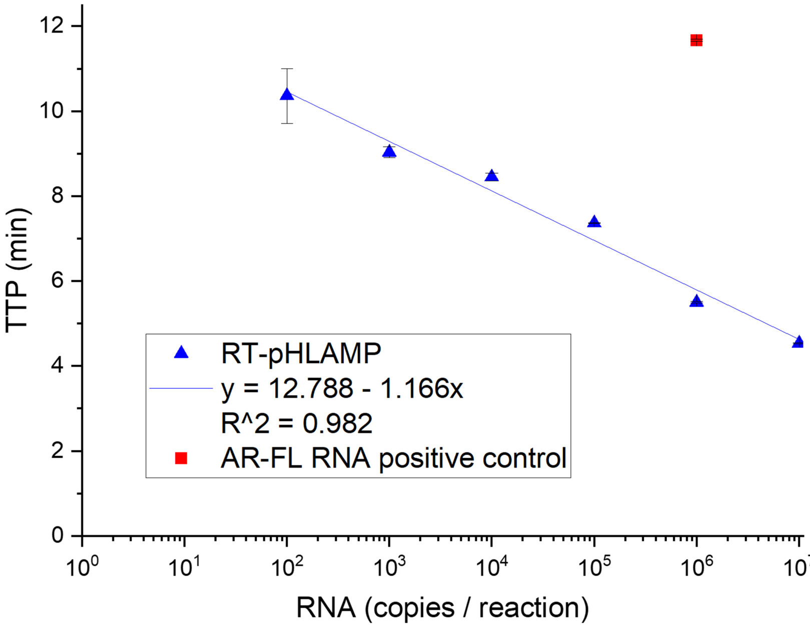

Figure 5: This graph shows the effect of synthetic AR-FL RNA presence on reducing the efficiency of the AR-V7 RT-pHLAMP assay. A positive control for AR-FL was included to confirm the presence of the RNA in these assays (shown in red).

#### 5 YAP1 RT-pHLAMP and serum reaction

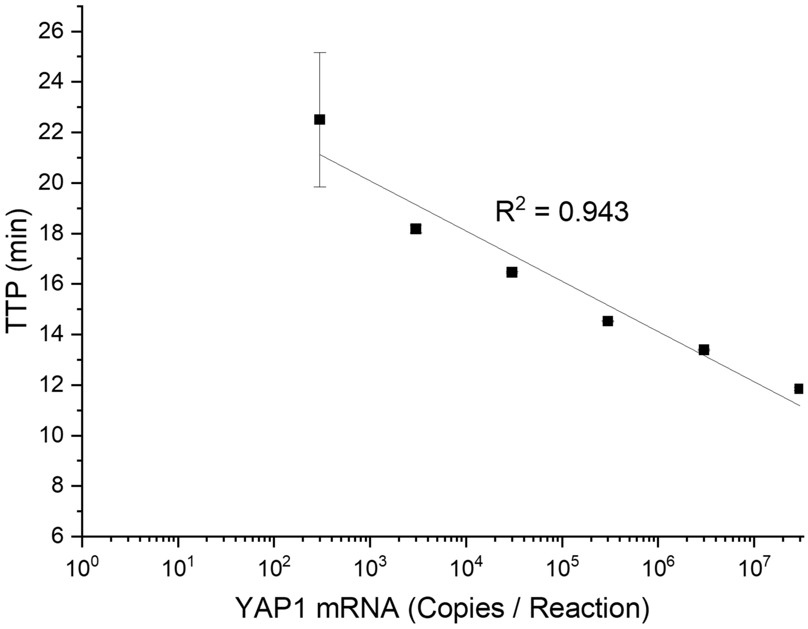

Figure 6: The standard curve for the RT-pHLAMP assay containing serum. RNA samples were diluted in mixed male serum before the assay was subsequently run.

### 6 qPCR data for AR-V7 mRNA in PCa cell lines

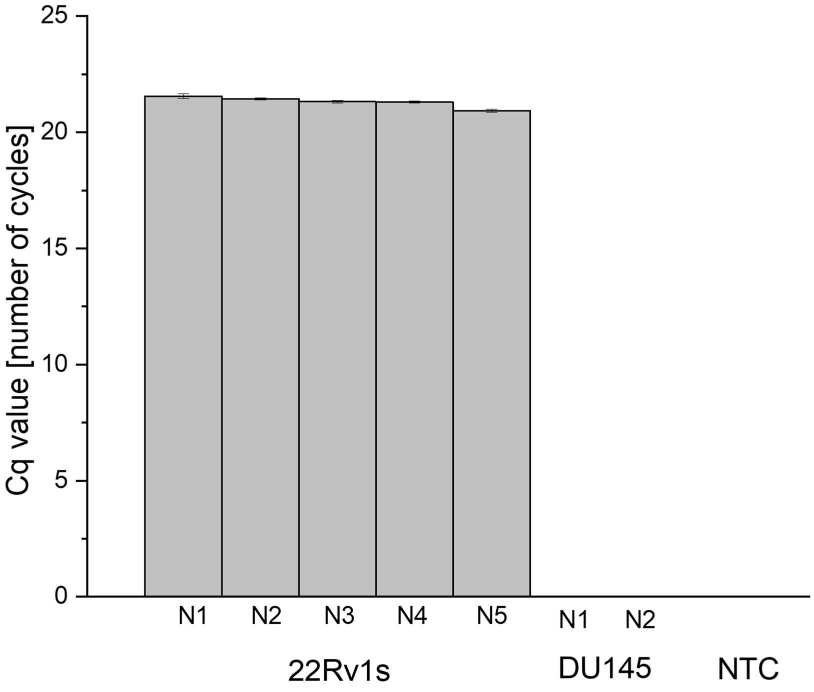

Figure 7: This shows the qPCR data in two prostate cancer cell lines for the presence of AR-V7. This assay was detecting cDNA which was reverse transcribed from extracted RNA from these cell lines. The same extracted RNA was used in the RT-pHLAMP reactions and Lab-on-Chip reactions in the main manuscript.

#### 7 qPCR data for YAP1 mRNA in PCa cell lines

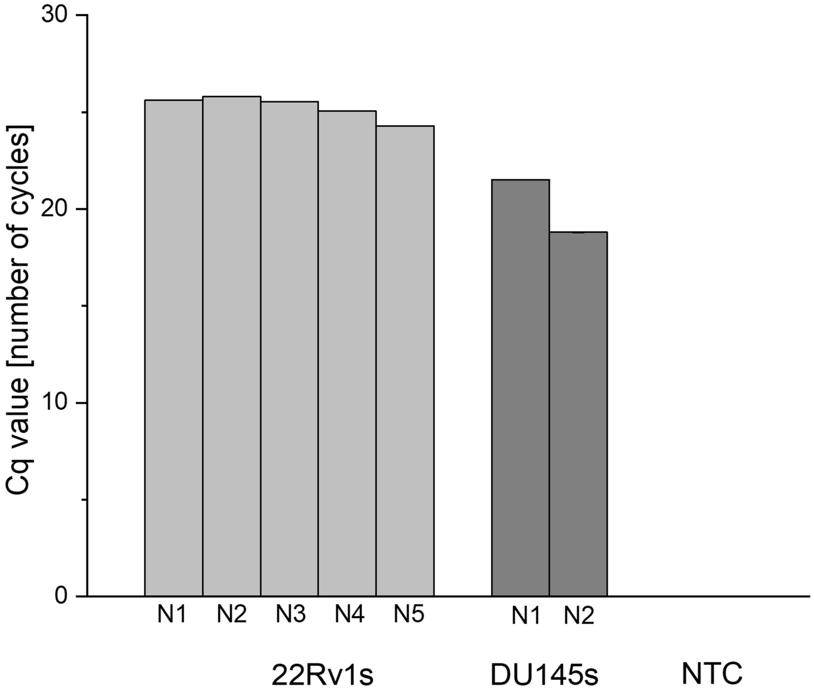

Figure 8: This shows the qPCR data in two prostate cancer cell lines for the presence of YAP1. This assay was detecting cDNA which was reverse transcribed from extracted RNA from these cell lines. The same extracted RNA was used in the RT-pHLAMP reactions and Lab-on-Chip reactions in the main manuscript.

#### 8 qPCR primer sequences

YAP1 forward primer 5'- GCACCTCTGTGTTTTAAGGGTCT - 3'
YAP1 reverse primer 5' - CAACTTTTGCCCTCCTCCAA - 3'
AR-V7 forward primer 5' - GACTCTGGGAGAAAAATTCCG - 3'
AR-V7 reverse primer 5' - CTCCAGACTATCCACTAGAG - 3'

### 9 Cq value determination

Lightcycler 96® (Roche diagnostics) devices utilise a predefined threshold fluorescence to calculate Cq values for SYBR green and SYBR green- like dyes. The predefined threshold for fluorescence for these fluorophores is 0.2.

### 10 Materials and Methods

#### 10.1 Synthesis of synthetic RNA targets

RNA fragments of AR-V7 and YAP1 sequences were synthesized from DNA gblocks (Integrated DNA Technologies) utilising the HiScribe™ T7 Quick High Yield RNA Synthesis Kit (NEB) according to the manufacturer's instructions including the DNase step. Stock concentrations were maintained at 3x$10^{10}$ copies per $\mu$L and stored at -80 $^o$C in preparation for experiments.

#### 10.2 RT-qLAMP experiments

All reactions were completed in triplicate. Each 10 $\mu$L experiment contained: 1 $\mu$L 10x isothermal buffer (New England Biolabs (NEB)), 0,6 $\mu$L $MgSO_4$ (100 mM stock), 1.4 $\mu$L dNTPs (10 mM stock of each nucleotide), 0.6 $\mu$L BSA (20 mg / mL stock), 0.8 $\mu$L betaine (5 M stock), 0.25 $\mu$L SYTO 9 green (20 $\mu$L stock), 0.25 $\mu$L NaOH (0.2 M stock), 0.042 $\mu$L Bst 2.0 DNA polymerase (120,000 U / ml stock, NEB), 0.1 Ribolock RNAse Inhibitor $\mu$L (40 U / $\mu$L stock, ThermoFisher), 0.3 $\mu$L Warmstart® RTx reverse transcriptase (15,000 U / ml, NEB), 1 $\mu$L 10x LAMP primer mix (20 $\mu$M FIP and BIP, 10 $\mu$M LB and LF, 2.5 $\mu$M F3 and B3), 1 $\mu$L RNA sample and the remaining solution was topped up to 10 $\mu$L with nuclease-free water. Reactions were conducted at 63$^o$C for 35 min. One melting curve from 63 $^o$C to 97 $^o$C was conducted to confirm the specific amplification of the reaction at a ramp of 0.2 $^o$C / s. Reactions were conducted with a Lightcycler® 96 instrument (Roche Diagnostics) in 96 well plates.

#### 10.3 RT-pHLAMP experiments

All reactions were completed in triplicate. Each 10 $\mu$L experiment contained: 1 $\mu$L customized isothermal buffer, 0.5 $\mu$L $MgSO_4$ (100 mM stock), 1.4 $\mu$L dNTPs (10 mM stock of each nucleotide), 0.6 $\mu$L BSA (20 mg / ml stock), 0.25 $\mu$L SYTO 9 green (20 $\mu$M stock), 0.25 $\mu$L NaOH (0.2 M stock), 0.042 $\mu$L Bst 2.0 Warmstart® DNA polymerase (120,000 U / mL stock, NEB), 0.3 $\mu$L Warmstart ® RTx reverse transcriptase (15,000 U / ml stock, NEB), 1 $\mu$L 10x LAMP primer mix (20 $\mu$M FIP and BIP, 10 $\mu$M LB and LF, 2.5 $\mu$M F3 and B3), 1 $\mu$L RNA sample and the remaining solution was topped up to 10 $\mu$L with nuclease-free water. For serum experiments, the RNA sample was diluted in mixed male serum (Sigma Aldrich) and 1 $\mu$L of that solution was added to the reaction. Reactions were conducted at 63$^o$C for 35 min. One melting curve from 63 $^oC$ to 97 $^oC$ was conducted to confirm the specific amplification of the reaction at a ramp of 0.2 $^oC$ / s. Reactions were conducted with a Lightcycler ® 96 instrument (Roche Diagnostics) in 96 well plates. Reactions were scaled up to either 12 $\mu$L or 20 $\mu$L reactions for implementation onto the LoC device, proportions of each reagent were kept the same.

#### 10.4 RT-qPCR experiments

All reactions were completed in triplicate. RT-qPCR reactions were completed in two steps. 50 ng mRNA samples were initially converted to cDNA with a RevertAid First Strand cDNA synthesis kit (ThermoFisher Scientific) as per the manufacturer's instructions including the optional step for GC rich regions. cDNA was used immediately for qPCR assays. qPCR experiments were conducted in 10 $\mu$L quantities and contained the following: 5 $\mu$L Fast SYBR® Green Master Mix (Applied Biosystems), 2 $\mu$L cDNA sample, 0.5 $\mu$L forward primer (250nM, 5 $\mu$M stock), 0.5 $\mu$L reverse primer (250nM, 5 $\mu$M stock), Nuclease-free water was added to make the reaction volume up to 10 $\mu$L. Reactions were aliquoted into a 96 well plate for analysis with a StepOnePlus™Real-Time PCR system (Applied Biosystems). Reactions were initially heated to 95 $^oC$ for 20 s. The cycling stage including heating at 95 $^oC$ for 3 s followed by 60 $^oC$ for 30 s. The cycling stage was repeated for 40 cycles. Melting curves were conducted with heating to 95 $^oC$ for 15 s followed by 60 $^oC$ for 1 min.

$$v = \frac{(\frac{s_1^2}{N_1} + \frac{s_2^2}{N_2})^2}{\frac{(\frac{s_1^2}{N_1})^2}{N_1 - 1} + \frac{(\frac{s_2^2}{N_2})^2}{N_2 - 1}} \tag{1}$$

The equation for $t$ value for the Welch's $t$-test of unequal variance is shown below (Equation **(2)**), where $\bar{x_1}$ and $\bar{x_2}$ are the mean values of the two datasets.

$$t = \frac{\bar{x_1} - \bar{x_2}}{\sqrt{\frac{s_1^2}{N_1} + \frac{s_2^2}{N_2}}} \tag{2}$$

The null hypothesis was rejected when $p < .05$.
